# Olfactory object recognition based on fine-scale stimulus timing in *Drosophila*

**DOI:** 10.1101/418632

**Authors:** Aarti Sehdev, Yunusa G. Mohammed, Tilman Triphan, Paul Szyszka

## Abstract

Odorants of behaviorally relevant objects (e.g., food sources) intermingle with those from other sources. Therefore, to sniff out whether an odor source is good or bad – without actually visiting it – animals first need to segregate the odorants from different sources. To do so, animals could use temporal cues, since odorants from one source exhibit correlated fluctuations, while odorants from different sources are less correlated. However, it remains unclear whether animals can rely solely on temporal cues for odor source segregation. Here we show that 1) flies can use a few milliseconds differences in odorant arrival to segregate a target odorant from a binary mixture, 2) segregation does not improve when the target odorant arrives first, and 3) segregation works for odorants with innate, as well as learned valences. These properties of odor segregation parallel those of concurrent sound segregation and figure-ground segregation by onset asynchrony in humans.

## INTRODUCTION

A natural scene is comprised of primary stimulus features, such as the spectral reflectance, intensity and movement of objects. In addition, it consists higher-order stimulus features that reflect the spatial and temporal coherence of those stimuli that belong to the same object (e.g., the correlated movements of a person’s body parts that allow us to segregate the person from the crowd). The mechanisms of how sensory systems use higher-order stimulus features for object recognition are well understood in vision [1] and audition [2], but not in olfaction. Olfaction research has mainly focused on primary stimulus features, such as chemical identity, concentration and dynamics of olfactory stimuli [3,4], yet it is still unknown how the olfactory system processes higher-order stimulus features that underlie olfactory object recognition.

Olfactory object recognition involves recognizing whether intermingling odorants originate from the same or different sources [5]. The capability to segregate odor sources is behaviorally relevant. For example, it allows animals to ignore spoiled food (food and detrimental odorants originate from the same source) and to find good food in a patch of spoiled food (food and detrimental odorants originate from different sources) without actually visiting the source.

Odor source segregation can be achieved from afar by analyzing the spatial distribution of odorants in a plume. This is because the different odorants from a single source form plumes with stable odorant concentration proportions (homogeneous plumes), while odorants from different sources form plumes with variable odorant concentration proportions (heterogeneous plumes) [5,6]. Correspondingly, plume heterogeneity enables animals to segregate odor sources (slugs: [7], insects: [8–11], crabs: [12]). But how do they do it? An animal could use spatial sampling to detect the spatial heterogeneity of odorant concentrations by comparing odorant stimuli along or between their olfactory organs. This strategy is plausible for animals with long olfactory tentacles (slugs), antennae (insects) and antennules (crabs), but this strategy might not work for animals with small and narrow olfactory organs, such as fruit flies, because they lack spatial resolution. Alternatively, an animal could use temporal sampling to detect time differences in odorant arrival for odor source segregation, as odorants from a single source exhibit more correlated fluctuations than odorants from different sources [5,6]. The latter strategy might be the only one available for small animals, such as fruit flies. This affords the possibility of using the fruit fly to investigate how temporal cues can be used for odor-object segregation.

The neural mechanism by which a heterogeneous odor plume is segmented into its constituent odor objects is unknown. Determining the causal relationship between behavioral odor source segregation and neural activity requires manipulating neural activity in identified neurons. As this is facilitated by the genetic tools available in the fruit fly *Drosophila melanogaster*, we here studied flies’ capability to use temporal stimulus cues for odor source segregation. We demonstrate that flies can use few milliseconds short differences in odorant arrival (referred to as onset asynchrony) to segregate odorants with different innate or learned valences. The flies’ high temporal precision of olfactory processing observed here lays the foundations for causal studies on the mechanisms of odor-background segregation and olfactory object recognition and implies a rapid and temporally precise mechanism for the encoding of olfactory objects.

## RESULTS

To ascertain whether flies can use stimulus onset asynchronies to segregate odorants in a plume, we used a free-flying behavioral paradigm in a wind tunnel to test flies’ preference to binary mixtures of attractive and aversive odorants with different onset times (Figure 1). We tracked flies’ flights in 3D and measured their odor preference using either odorants with innate or conditioned valence (original data are available in Data S1). The pairs of odorants with innate valence used were 2-butanone (BN) and butanal (BA), and BN and benzaldehyde (BZ). For the conditioned odorants, we used 2,3-butanedione (BD) and ethyl acetate (EA). We chose these odorants based on their innate valences measured in tethered flying flies [13], in which BN was attractive, whereas BA and BZ were aversive, and BD and EA were behaviorally neutral.

**Figure 1.**
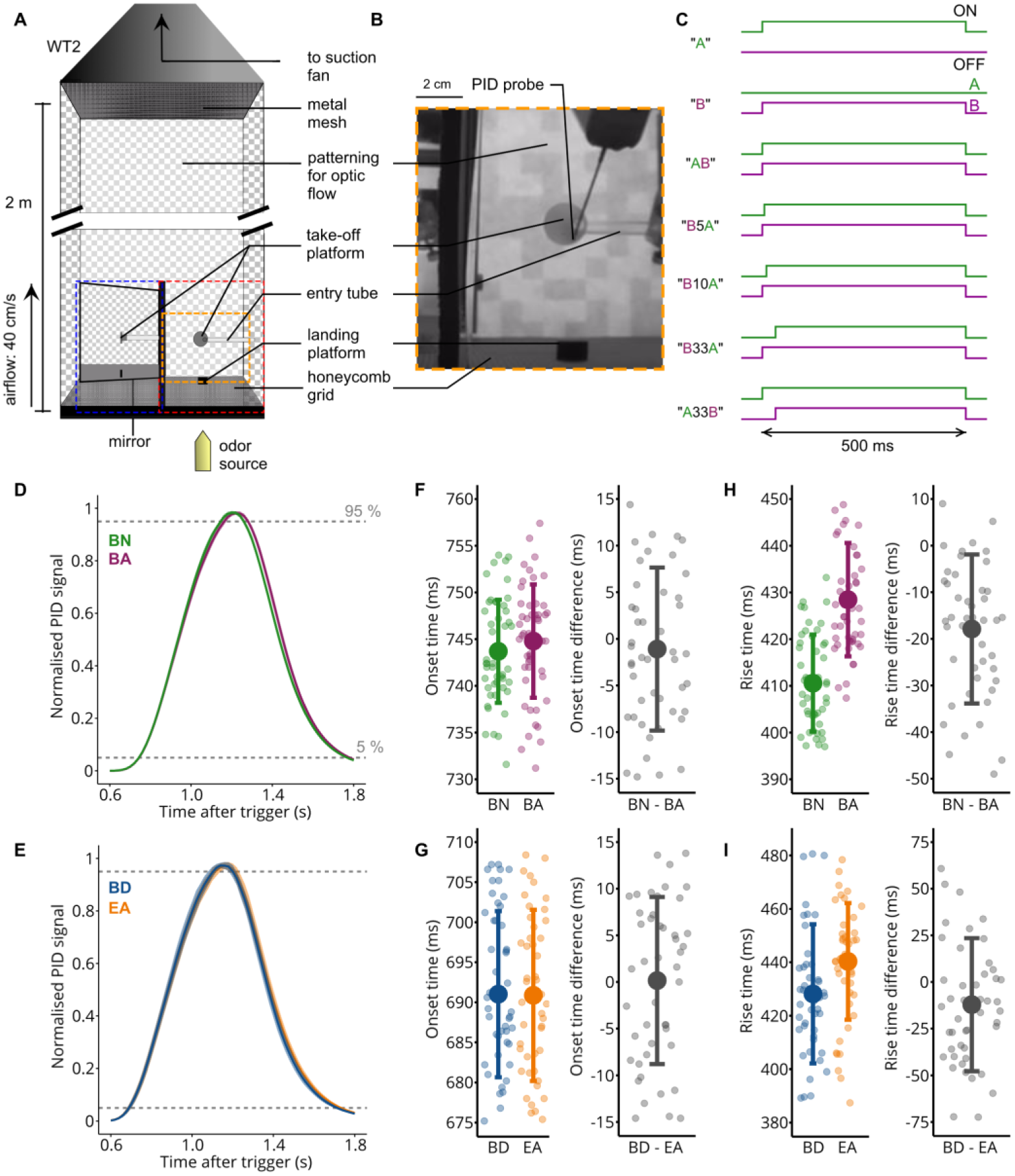
Delivering temporally precise olfactory stimuli in a wind tunnel. (**A**) Diagram of wind tunnel 2 (WT 2). Red and blue dashed boxes indicate the captured x-y and z-y planes respectively. The olfactory stimulator was placed outside of the wind tunnel to minimize turbulences. The orange box outlines in the diagram the image in (B). (**B**) The layout of WT 2, showing the position where the odorant concentrations were recorded using a PID. (**C**) Valve states for the different stimuli. The attractive odorant A and aversive odorant B are represented in green and magenta respectively. When asynchronous mixtures were presented, the first odorant was always given for 500 ms, and the following odorant with an onset delay. Both odorants had the same offset time. Pulses were repeated every 2 s. (**D**) PID recordings of pulsed stimuli for the odorant pair with innate valence 2- butanone (BN, green) and butanal (BA, magenta) (mean and SD over 50 pulses). Valves opened for 500 ms. Each PID signal was normalized to the maximum concentration reached. (**E**) Same as (D) for the odorant pair with conditioned valence 2,3-butanedione (BD, blue) and ethyl acetate (EA, orange), averaged over 50 pulses. (**F**) Left: Onset time (time taken to reach 5 % of maximum concentration after valve trigger) for BN and BA (mean and SD over 50 pulses). Individual points represent the onsets for each pulse. Right: Onset time difference between pairs of BN and BA pulses (mean and SD over 50 pulses). (**G**) Same as (F) for BD and EA. (**H**) Left: Rise time (time take to reach 95 % of maximum concentration from the 5% onset time) for BN and BA (mean and SD over 50 pulses). Individual points represent the rise times for each pulse. Right: Mean rise time difference between pairs of BN and BA pulses (mean and SD over 50 Pulses). (**I**) Same as (H) for BD and EA.

To mimic homogeneous odorant plumes from one source we presented both odorants as a synchronous mixture (no onset delay between odorants), and to mimic heterogeneous odorant plumes from different sources we presented both odorants as asynchronous mixtures (with 5 to 33 millisecond delays between odorant onsets) (Figure 1C). Note that we used different wind tunnels (WT 1 and WT 2) and different arrangements of the odor source and landing platforms in an attempt to optimize experimental conditions. However, we found no clear differences in flies’ performance, indicating that the results are robust, and do not depend on peculiar arrangements of the wind tunnels. To eliminate between-session variability, all data shown in a given plot were collected in parallel. Accordingly, data points should be compared within plots but not between plots.

### Tracking of temporally well-controlled odorant stimuli in the wind tunnel

We initially determined how reliable our stimulus delivery was over time by using a photoionization detector (PID) to record the stimulus dynamics of the different odorants used (Figure 1D - 1I, S1A-C). The inlet of the PID was placed at the surface of the take-off platform (Figure 1B). Each odorant was presented 50 times within its odorant pair BN and BA, BN and BZ, or BD and EA. The onset times (time it took from valve opening to reach 5 % of the maximum PID signal) were temporally precise across trials, with standard deviations ranging between 6 ms (BN, BA) and 10 ms (BD, EA) (Figure 1F, 1G and S1B).

The onset times were similar for all odorant pairs (BN/BA, BN: 744 ms ± 6 ms, BA: 745 ms ± 6 ms; BN/BZ, BN: 750 ms ± 7 ms, BZ: 756 ms ± 7 ms; BD/EA, BD: 691 ms ± 10 ms, EA: 691 ± 10 ms; mean ± SD). The rise times (time it took to reach from 5 % to 95 % of the maximum PID signal) were also similar for the odorant pair BN/BA (BN: 411 ± 10 ms, BA: 428 ± 12 ms; mean ± SD) and for the odorant pair BD/EA (BD: 428 ms ± 26 ms, EA: 440 ms ± 21 ms), but less similar for the odorant pair BN/BZ (BN: 400 ms ± 12 ms, BZ: 444 ms ± 9 ms) (Figure 1H, 1I and S1C). The differences in stimulus dynamics could be explained by the difference in the molecular mass between odorants, as stimulus dynamics get slower with increasing molecular mass (in g/mol, BN: 72; BA: 72; BD: 86; EA: 88; BZ: 106) [14,15].

To visualize how flies explored space based on the odorant experience, we projected their flight trajectories on a plane, and calculated the probability across flies to visit a particular pixel (visit probability, Figure 2 A). When presented with the innately attractive odorant BN, flies were more likely to fly towards the target (which was either the actual odor source or a black platform near the odor source, see Methods) compared to the innately aversive odorant BZ. To assess approach to the target, we counted the number of flies which reached halfway between the center of the take-off platform and the target (3.1 cm (117 pixels) for WT 1 and 2.7 cm (71 pixels) for WT 2) and calculated the approach probability by dividing this number by the total number of flies. Flies flew closer towards the target when stimulated with an attractive odorant than with an aversive odorant or a control air stimulus (Air) (*p(BN > BZ)* ≥ 0.999 in Figure 2B; *p(BN > BA)* = 0.962*, p(BN > Air)* ≥ 0.999 in Figure 2C; all statistical significances are given as Bayesian probabilities, see Methods) Figure. However, in contrast to previous studies [16–19], flies rarely landed at or near the target. This discrepancy might reflect the fact that, different to these previous studies, our odorant delivery device was outside the wind tunnel. Positioning the odor delivery outside the wind tunnel prevents turbulences which could provide localization cues for the fly to land. Rather, our wind tunnel setting mimics better an odor source at distance.

**Figure 2.**
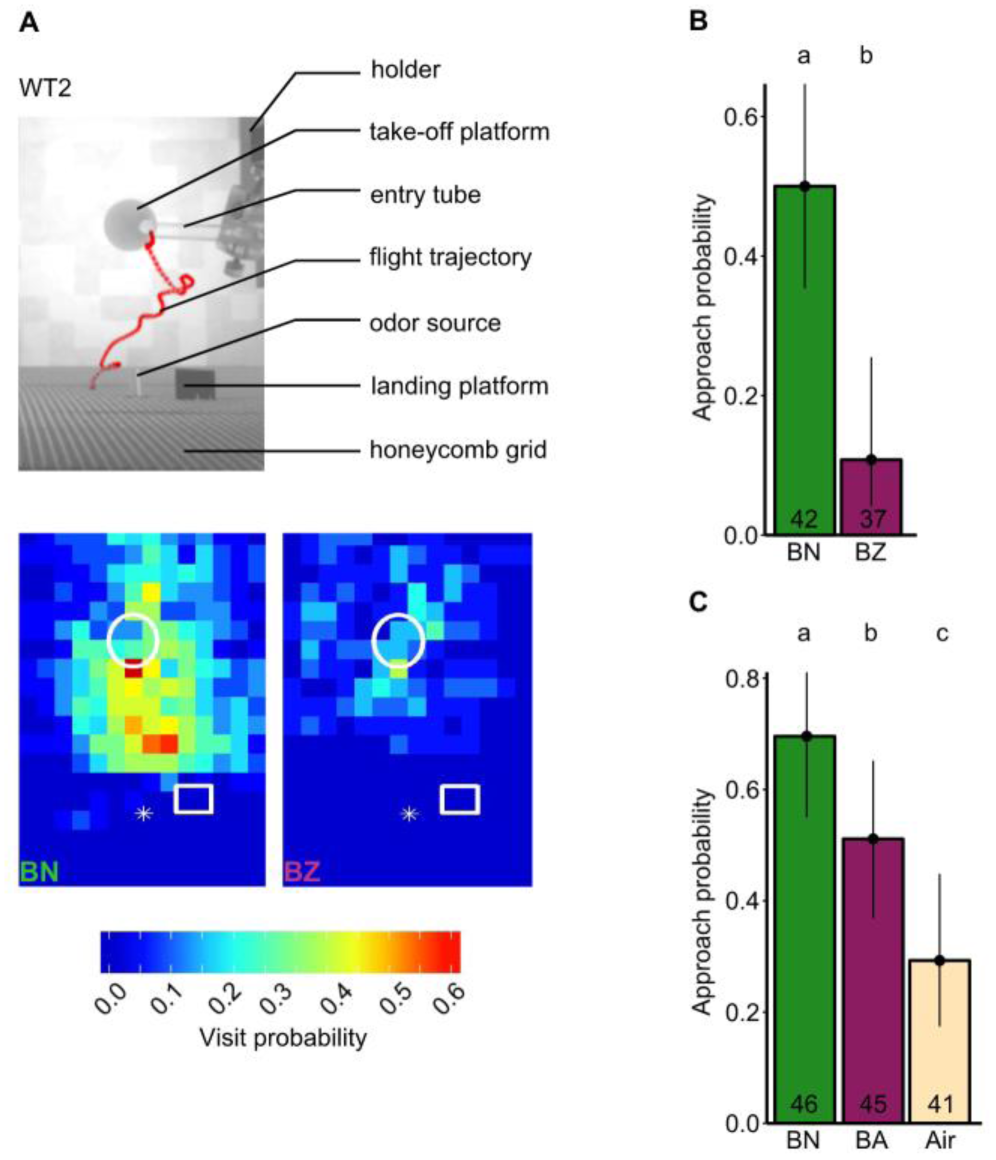
Odor tracking in the wind tunnel. See also Figure S1. (**A**) Top: Minimum intensity projection of a movie of a flying fly (red) in the wind tunnel during stimulation with BN. Bottom: Visit probability plot equivalent to top image for BN and BZ respectively (set 1). Each bin represents 20 x 20 pixels in the image, corresponding to 7.6 x 7.6 mm at the height of the landing platform. Each bin shows the mean binary value across flies. The take-off platform (white circle), landing platform (white rectangle) and odor source (white star) are indicated for position reference. n = 24 and 20 for BN and BZ respectively. (**B**) Approach probability to cross the half distance between take-off platform and landing platform for BN and BZ. Bars represent the fitted mean from the GLM. Error bars represent the 95 % credible intervals. The lower case letters represent significantly different responses for the different odorants; this applies throughout the figure. Numbers in bars indicate the number of flies; this applies throughout all figures. (**C**) Same as in B but for BN, butanal (BA) and a blank air stimulus (Air).

The percentage of flies that started flying ranged between 85 to 96 % for the attractive odorant BN, 68 to 84 % for the aversive odorants BZ or BA and 71 % for the blank air control (Table S1). The average latency to flight ranged from 10 – 27 s, corresponding to 5 to 13 odorant stimuli before taking off (Table S1, Figure S1H, S1J and S2D, S2E). There was no consistent connection between the valence and of an odorant and the latency to flight (e.g., latency to flight was longer for BZ than for BN (Figure S1H), but there was no difference for BN and BA (Figure S1J).

### Attraction towards asynchronous mixtures of odorants with differing innate valence

To test whether flies can detect stimulus onset asynchrony, we presented the attractive odorant BN (A) and the aversive odorant BA (B) either as single odorants, combined in a synchronous mixture (AB) or in asynchronous mixtures in which B preceded A by 33 ms (B33A) (Figure 3A). Note that we used the odorant pair BN/BA to test the effect of stimulus onset asynchrony rather than BN/BZ because the differences in stimulus dynamics between BN and BZ makes this odorant pair unsuitable for generating synchronous mixtures (Figure S1).

**Figure 3.**
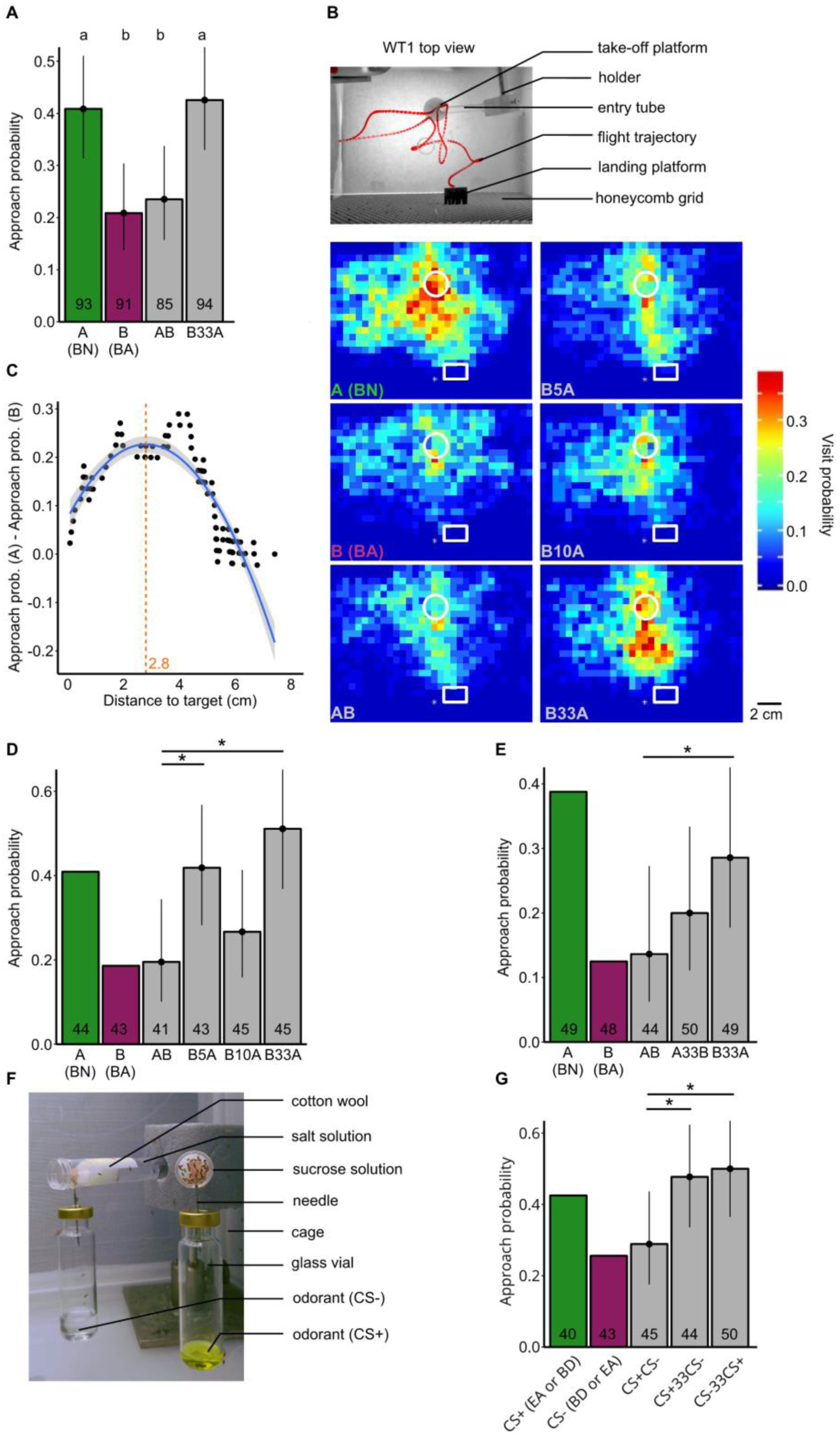
Stimulus onset asynchrony makes a mixture of odorants with differing valences more attractive. See also Figure S2 and S3.(**A**) Approach probability determined by the half distance-threshold for the single odorants BN (A), BA (B), their synchronous mixture (AB) and their asynchronous mixture (B33A). Bars represent the fitted value from the GLM. Error bars represent the 95 % credible intervals. The lower case letters represent significantly different responses to the odorant treatments (this dataset is pooled from experiments shown in (D) and (E)). (**B**) Top: Minimum intensity projection of a movie of a flying fly (red) in the wind tunnel during stimulation with BN (A). Bottom: Visit probability maps for A (BN) and B (BA) and the synchronous (AB) and asynchronous (B5A, B10A, B33A) mixtures. The take-off platform (white circle), landing platform (white rectangle) and odor source (white star) are indicated for position reference. n = 44, 43, 41, 43, 45 and 45 for A, B, AB, B5A, B10A and B33A respectively. (**C**) Thresholding method that uses the distance which separates flies’ approach probabilities for A and B best (maximized A-B difference threshold). Each point represents the proportion of A-stimulated flies that approached the target by the given minimum distance to the target minus the proportion of B-stimulated flies. The blue trend line was fitted using locally weighted scatterplot smoothing. The distance at the peak of the trend line was defined as threshold (orange dashed line and value). (**D**) Approach probability for odorant mixtures with different asynchronies (maximized A-B difference thresholding method). Stars represent significantly different responses between AB and the other mixtures. Since A and B are used to determine the threshold, they were not included in the statistical analysis and the observed rather than fitted means are shown. (**E**) Approach probability for odorant mixtures with different odorant orders (maximized A-B difference thresholding method). (**F**) Conditioning setup in which flies were left for autonomous differential conditioning. Flies can freely fly in the cage and enter the odorized tubes containing cotton wool soaked either with aversive salt solution or attractive sucrose solution. (**G**) Approach probability for odorant mixtures with conditioned valences (maximized A-B difference thresholding). Odorants BD and EA were used equally as often as the CS+ and CS-.

Flies showed a higher approach probability for A compared to B (*p(A > B*) = 0.998) or to the synchronous mixture AB (*p(A > AB)* = 0.993) (Figure 3A). Moreover, flies showed a higher approach probability for the asynchronous mixture B33A compared to synchronous mixture AB (*p(B33A > AB)* = 0.996) or to the aversive odorant B (*p(B33A > B)* ≥ 0.999). This shows that flies perceive the synchronous mixture AB and the asynchronous mixture B33A differently, with the onset asynchrony making the mixture more attractive.

To test whether flies are sensitive for shorter onset asynchronies we applied synchronous and asynchronous mixtures which started with B and with onset times differing by 5, 10 or 33 ms (B5A, B10A, B33A) (Figure 3B). Flies presented with A showed more activity in general, along with a higher visit probability near the target compared with flies presented with B. Flies showed a similar visit probability map for the synchronous mixture AB as for B. However, when stimulated with the asynchronous mixtures B33A or B5A – but not B10A, flies showed more activity near the target compared to AB and B.

To make the quantification of flies’ approach behavior less arbitrary and to account for the fact that flies distributed differently in the two different wind tunnels and experimental sets, we calculated an approach area that segregated flies’ approach probabilities for the attractive odorant A and the aversive odorant B the most (Figure 3C, Figure S1-S3). We determined this area for each experimental set separately (see Methods). Note that this method maximizes the differences in approach probability between odorants A and B by design. Therefore, we refrain from comparing flies’ approach probabilities for A or B and restrict the comparisons to the mixtures.

The flies’ responses to the mixtures depended on the timing between B and A (Figure 3D). For onset asynchronies of 5 ms (B5A) and 33 ms (B33A), flies were attracted to the target and scored a higher approach probability than for the synchronous mixture AB (*p(B5A > AB)* = 0.984, *p(B33A > AB)* = 0.998), similar to that of A alone. However, for the onset asynchrony of 10 ms (B10A), flies’ approach probability was not different to the approach probability for AB (*p(B10A > AB)* = 0.783). While we have no good explanation for this non-monotonic relationship between onset asynchrony and behavioral performance, this effect is reminiscent of visual masking in humans, in which masking efficiency also depends non-monotonically on the asynchrony between target and masking stimulus [20].

Next, we wanted to discern whether the order in which odorants are presented in a mixture affects how a fly perceives the mixture. We used the same paradigm and odorants as before and stimulated flies with the synchronous mixture AB, the asynchronous mixture A33B (A precedes B) and B33A (B precedes A) (Figure 3E and S2). In this paradigm, flies showed a lower approach probability to the synchronous mixture AB than to the asynchronous mixture B33A (*p(B33A > AB)* = 0.957)), confirming our previous result that B33A is perceived differently to AB, and is perceived by the fly as more attractive. However, the approach probability for the asynchronous mixture A33B was not significantly different to the approach probability for AB *(p(A33B > AB)* = 0.793)), indicating that the two asynchronous mixtures A33B and B33A may have been also perceived differently.

These data show that flies can discriminate between the synchronous mixture AB and asynchronous mixtures B5A and B33A, supporting the hypothesis that flies can use stimulus onset asynchrony to segregate the attractive component A from the mixture of A and B even if they never encountered A alone (in B5A and B33A, B started before A and A ended at the same time as B). In contrast, the similar low approach probabilities for the aversive odorant B and the synchronous mixture AB is consistent with the hypothesis that flies perceive AB as coming from one source.

### Attraction towards asynchronous mixtures of odorants with differing learned valence

Finally, we wanted to determine whether flies’ capability to discriminate between synchronous and asynchronous mixtures only works for odorants with differing innate valence, or whether it also works for odorants with differing learned valences. To address this question, we used an autonomous differential conditioning paradigm and paired one odorant (positively conditioned stimulus, CS+) with a 1M sucrose solution and another odorant (negatively conditioned stimulus, CS-) with a saturated NaCl solution (Figure 3F). We used the odorants EA and BD equally often for CS+ and CS-. This procedure eliminates all non-associative effects of the conditioning procedure (e.g., sensitization), which would also change flies’ responsiveness [21]. Thus CS+ and CS-only differ with regards to the learned valences, devoid of odorant-specific innate valences. Also in this experiment, flies discriminated between synchronous and asynchronous mixtures, and showed lower approach probabilities to the synchronous mixture of the CS+ and the CS-(CS+CS-) than to the asynchronous mixture CS+33CS- or CS-33CS+ (*p(CS+33CS- > CS+CS-)* = 0.965, *p(CS-33CS+ > CS+CS-)* = 0.981) (Figure 3G and S3). Together, these findings support the hypothesis that flies can use stimulus onset asynchrony to segregate odorants with both learned and innate valences from mixtures.

## DISCUSSION

We asked whether *Drosophila* can use stimulus onset asynchrony to segregate an attractive target odorant from a mixture with a less attractive odorant. We found that flies show stronger attraction to asynchronous mixtures (mimicking two odorant sources) than to synchronous mixtures (mimicking one source). These results demonstrate a hitherto unknown temporal precision in olfaction and show that the fly’s olfactory system can use stimulus onset asynchrony for olfactory object recognition and odor-background segregation.

### Odor-source segregation

Previous studies showed that animals perceive different odorants from the same source as one object while they perceive odorants from different sources as separate objects. In a pioneering study, Hopfield and Gelperin [7] aversively conditioned slugs to the mixture of two food odors, A and B. When A and B were homogeneously mixed during conditioning (to mimic one odor source), slugs afterwards responded aversively to the homogenous mixture AB but not to the single odorants A and B. However, when A and B were heterogeneously mixed during conditioning (to mimic different sources), slugs afterwards responded aversively also to A and B alone. This suggests that slugs perceived the homogenous mixture AB as different from A or B, while they perceived the heterogeneous mixture as distinct odor objects A and B. Similarly, several arthropods species can segregate attractive from aversive odorants depending on whether both are released from the same source (forming a homogeneous mixture) or from different sources (forming a heterogeneous mixture) [8–12, 22]).

In the above studies, animals could have achieved odor source segregation by detecting the heterogeneous distribution of odorants through a spatially heterogeneous activation across or within their olfactory organs, or they may have recognized the single odorants during bouts of their pure, unmixed presence. Compared to the above animals, *Drosophila* has tiny olfactory organs. Therefore, in lack of spatial resolution, *Drosophila* might use temporal rather than spatial stimulus cues for odor source segregation. Our data suggest that the critical asynchrony for odor segregation is in the range between 5 to 33 ms. Importantly, odor segregation works when the target odorant is never encountered alone (in BΔtA, the target odorant A is always mixed with B, because A starts after and ends with B). These properties of odor segregation in *Drosophila* parallel those of concurrent sound segregation [23,24] and figure-ground segregation [25,26] in humans in which the critical asynchrony for segregation is around 30 ms and segregation is no better when the target is presented first.

### Mechanisms of odor source segregation

The odor-source segregation paradigms, that were used in previous studies and in the present study, were odor recognition tasks in which the odorants either had innate valences [8,9,12,22] or learned valences [7,10,11], and it is unknown whether animals can segregate mixtures of novel odorants that have no innate or learned valence. Thus, in previous studies and our own study, to recognize the odorants A and B, the olfactory system has to match the odor-evoked neural activity patterns to a neural template of A and B. The neural templates could have developed through evolution (e.g., odorants with innate valence activate specific, valence-encoding neurons in the lateral horn [27–30]), or by associative learning (odorants with learned valence activate specific, valence-encoding neurons in the mushroom body [31–33]).

Flies’ capability to segregate two mixed odorants A and B based on a few milliseconds onset asynchrony poses temporal constraints on the neural code for odors. The computations that the olfactory system could use to perform odorant segregation are coupled to how the animal perceives the single odorants and their mixtures. As we do not know what the flies actually smell, but we can measure their attraction towards the odorants, we can only speculate about the perceptual differences between synchronous and asynchronous mixtures. In the following we shall discuss two alternative mechanisms of odor source-segregation based on temporal stimulus cues.

### Shift from synthetic to analytic mixture processing?

Flies could perceive the synchronous mixture AB synthetically such that information about the components A and B is lost (AB ≠ A + B), and they could perceive the asynchronous mixture A∆tB analytically such that information about A and B is preserved (A∆tB = A + B).

Behavioral experiments in honey bees provide support for synthetic processing of synchronous mixtures: when conditioned to an odorant mixture, bees show lower response probabilities for the individual components than for the conditioned mixture [34], and bees can solve biconditional discrimination [35] and negative patterning tasks [36] which both require synthetic mixture processing. Physiological experiments also indicate that synchronous mixtures are processed synthetically, while asynchronous mixtures are processed more analytically: mixing of multiple odorants alters the neuronal response patterns across olfactory receptor neurons and second-order olfactory neurons (projection neurons) such that component information gets partly lost [37– 41]. In contrast, the responses of projection neurons to asynchronous mixtures partly match those evoked by the individual odorants, with the first arriving odorant often dominating the response pattern [11,37,42,43]. However, such dominance of the first arriving odorant occurred neither in behavioral experiment in honey bees [10] nor in flies (this study). We therefore conclude that an asynchrony-induced shift from synthetic to a more analytic mixture representation cannot fully explain the behavioral odor source segregation observed in flies.

### Analytical mixture processing and parallel encoding of source separation?

Alternatively, flies could perceive both synchronous and asynchronous mixtures analytically (AB = A +B and A∆tB = A + B), and the information that odorant A and B belong to the same or to different sources could be encoded in parallel by the timing between A- and B-activated identity- or valence-encoding neurons.

Although there is evidence for synthetic processing of synchronous mixtures in insects [34–36], there is also evidence for analytical mixture processing: when honey bees are trained to respond to a multi-odorant mixture and afterwards are tested with the single odorants, they respond to most of the odorants [44,45]. Analytic mixture processing has also been demonstrated in blocking experiments, in which previous conditioning to odorant A reduces (or blocks) conditioning to B during training with AB, because A already predicts the reward [46]. Experiments in *Drosophila* provide further evidence for analytical mixture processing, as flies’ responses to the synchronous mixture of two odorants with opposing valences A and B add up linearly [13,47]. Moreover, *Drosophila* fails in biconditional discrimination or negative patterning tasks, which require synthetic mixture processing, suggesting that *Drosophila* processes mixtures analytically [48].

In accordance with these behavioral indications of analytic mixture perception, neuronal response properties would support analytical mixtures processing: even though mixtures suppress the response strength of olfactory neurons, those neurons that respond strongly to the components generally also respond strongly to the mixture [11,37–39,43]. Moreover, *Drosophila* Kenyon cell responses to a mixture AB resemble the superposition of their responses to the single components A and B [49]. Therefore, the neuronal representations of both synchronous and asynchronous mixtures likely contain sufficient odorant component information to allow for analytic mixture processing.

Whether or not two odorants A and B originate from one or two sources could be detected by coincidence-detecting neurons that receive input from valence-encoding neurons of the lateral horn (for odorants with innate valences; [27–30]) or from output neurons of the mushroom body (for odorants with learned valences; [31–33]). Those coincidence-detecting neurons would respond to synchronous input from the A- and B-activated valence-encoding neurons (A and B come from one source) but not to asynchronous input (A and B come from different sources). Coincidence detection could, for example, be mediated by NMDA glutamate receptors [50]. The existence of glutamatergic neurons and NMDA receptors in both the lateral horn and in the mushroom body [51], and of glutamatergic valence-encoding mushroom body output neurons in *Drosophila* [32,52], is consistent with this hypothetical mechanism.

Detecting asynchronies of a few milliseconds between the arrival of odorant A and B requires temporally precise encoding of odorant onsets – a requirement that appears to be fulfilled by insect olfactory receptor neurons [53– 55]. In particular, *Drosophila* olfactory receptor neurons respond to odorants rapidly (with first spike latencies down to 3 ms) and across neurons of the same type, the standard deviation of the first spike latencies can be as low as 0.2 ms [56]. This high temporal precision of first odorant-evoked spikes across olfactory receptor neurons would allow a rapid, spike timing-based coding scheme for odorant onset and identity [14,56,57], which could underlie flies’ capability to segregate odorants based on onset asynchrony.

## Acknowledgements

We thank Stefanie Neupert for advice on the statistics and comments on the manuscript, C. Giovanni Galiza, Thomas Nowotny and Mario Pannunzi for comments on the manuscript and Cansu Tafrali for help with the experiments. This project was funded by the Human Frontier Science Program (RGP0053/2015 to PS).

## Author contributions

PS conceptualized and designed the study. YMG and AS performed the data collection. TT wrote the video processing script and provided expertise on the analysis. YMG and AS performed the video processing. AS performed the statistical analysis. PS, YMG and AS wrote and edited the manuscript. PS supervised the study.

## Data and materials availability

All data is either supplied in this paper or in the Supplementary Information, or can be requested directly from the authors.

## METHODS

### Animals

Wild-type Canton S *Drosophila melanogaster* were reared on standard medium (100 mL contain 7.1 g cornmeal, 6.7 g fructose; 2.4 g dry yeast, 2.1 g sugar beet syrup, 0.7 g agar, 0.61 ml propionic acid, and 0.282 g ethyl paraben) under a 12:12 hours light:dark cycle (light from 09:00 to 21:00), at 25 °C and 60% relative humidity. All flies used in the experiments were female, aged between four and eight days old.

### Wind tunnel

We carried out experiments in two wind tunnels, referred to here as wind tunnel 1 (WT 1, data shown in Figure 3D) and wind tunnel 2 (WT 2 data shown in all other figures). We filmed each experiment using Raspberry Pi cameras (Raspberry Pi Camera Module v2; Raspberry Pi 3 model B) for 2 or 3 minutes with a resolution of 640 x 480 pixels and 90 frames s^-1^; the first 10 seconds of flight duration was used for the analysis.

Both wind tunnels were constructed from clear Plexiglas. The inner side walls and floor were covered by a random checker board pattern (grey on white paper). The dimension of WT 1 was 1.2 m x 0.19 m x 0.19 m and of WT 2 was 2 m x 0.40 m x 0.40 m. The exhaust took in room air (28 °C, 60 % relative humidity) through the tunnel and removed it from the setup building via a ventilation shaft. An aluminum honeycomb grid (hole diameter x length: 0.53 cm x 3 cm, WT 1; 0.32 cm x 9.7 cm, WT 2) at the inlet and a grid at the outlet of the tunnel created a laminar flow throughout. The wind speed was 0.4 m s^-1^. We injected odorants into the inlet of the wind tunnel with an olfactory stimulator [15]. The outlet of the olfactory stimulator was 1 cm in diameter and was placed just outside of the honey comb grid, creating a laminar odorant plume within the tunnel. Flies entered the tunnel through a glass tube that was connected to a take-off platform whose center was 7.5 cm (WT 1) or 6 cm (WT 2) downstream from the inner side of the honeycomb grid. We also placed a black platform near the odor source, as recent studies have demonstrated that *Drosophila* stimulated by an attractive odorant approach dark spots [18,19]. In WT 1 we used two cameras to film the flies. One camera was placed above the wind tunnel to capture the x-y plane of movement, whereas the other was placed at the side of the wind tunnel (90° to the other camera), thus capturing the movement of the fly within the z-y plane. The volume filmed measured 17.3 cm x 17.3 cm x 13.0 cm (x, y, z). In WT 2 we used a single camera placed above the wind tunnel to record the fly trajectories in the x-y plane. In order to capture the z-y plane of the flight track, we positioned a mirror at a 45° angle to the camera inside of the wind tunnel. The volume filmed measured 13.7 cm x 10.3 cm x 9.5 cm (x, y, z). Both wind tunnels were illuminated with indirect, homogeneous, white light with a color temperature of 6500 K (WT 1: compact fluorescent light, tageslichtlampe24.de; WT 2: LEDs, led-konzept.de). Additionally, we used 830 nm backlight illumination to get contrast-rich images of the flies.

### Odorant delivery

Odorants were delivered into the wind tunnels using a custom-made multichannel olfactory stimulator [15]. All odorants were supplied by Sigma Aldrich. Pure odorants were stored in 20 ml glass vials (Schmidlin) sealed with a Teflon septum. The cross section of the odorant surface was 3.1 cm^2^. The headspace of odorized air was permanently drawn into the air dilution system using flowmeters (112-02GL, Analyt-MTC) and an electronic pressure control (35898; Analyt-MTC). The stimulator had three channels: one for each odorant and one for blank air. The odorant vials were constantly flushed with clean air throughout the experiment, so that the headspace concentration reached a steady state of odorant evaporation into the air and odorant removal by the air flush. Note that due to the permanent air stream the headspace odorant concentration never saturated. The total flow per odorant channel was always 300 ml min^-1^. In WT 1, BN was released at 50 ml min^-1^ and added to 250 ml min^-1^ air, and BA was released at 30 ml min^-1^ and added to 270 ml min^-1^ air (experiments in Figure 3). In WT 2, BN, BA and BZ were released at 50 ml min^-1^ and were added to 250 ml min^-1^ air (experiments in Figures 2 and 3). For the conditioned odorants we used the PID to determine the head space concentrations in the conditioning tubes (see below) by moving the PID needle rapidly into the conditioning tubes to prevent dilution in odorant concentration due to air suction of the PID. These concentrations from the conditioning paradigm were then adjusted in the odor delivery device by measuring the odorant concentration just above the take-off platform with the PID. EA was released at 4 ml min^-1^ and added to 296 ml min^-1^ air, and BD was released at 1.84 ml min^-1^ and added to 298.16 ml min^-1^ air (experiments in Figure 3).

The two odorant channels and a blank channel (each with an airstream of 300 ml min^-1^) were combined and injected into a carrier air stream of 410 ml min^-1^ and, resulting in a total air flow at the outlet of the stimulator of 1.31 L min^-1^, and a wind speed of 0.4 ms^-1^.

Stimuli were presented either as single odorants (either A or B), as a synchronous mixture of odorants presented simultaneously (AB) or as an asynchronous mixture, with different time delays between the release of the odorants. In B∆tA, B starts before A, with ∆t being either 5 ms, 10 ms or 33 ms. In A∆tB, A starts before B, with ∆t being 33 ms (Figure 1C). Note that the trailing odorant ended at the same time as the preceding odorant. Stimuli were delivered in odorant pulses of 500 ms, and the interstimulus interval was 2 s. To exclude that differences in flies’ approach behavior towards the asynchronous and synchronous mixture reflected responses to mechanical cues produced by valve switching, we applied the single odorants together with a 33 ms delayed blank stimulus (both stimuli ended at the same time).

During experiments, all odorants were removed from the wind tunnel via an exhaust into the outside atmosphere. Between experiments using different odorants, the stimulator valves were flushed out over night to remove any residual odorant. Valves were controlled by compact RIO systems equipped with digital I/O modules Ni-9403 and odorant delivery was controlled by software written by Stefanie Neupert in LabVIEW 2011 SP1 (National Instruments).

### Experimental protocol for odorants with innate valence

Day 1: Between 13:00 and 16:00, approximately 100 adult flies were removed from standard corn meal agar food and were subjected to food and water starvation for 24 hours in a cage (30×30×30 cm, BugDorm-1, BugDorm) that allowed them to move around freely, in a room with an approximate relative humidity of 60%, a temperature of 25 - 28 °C and 12 hour daylight cycle.

Day 2: Between 15:00 and 20:00, individual, flying female flies were removed from the cage and placed into a PVC tube through which they could walk freely to enter the wind tunnel and reach the take-off platform. Once the fly reached the take-off platform, odorant stimulation started. Each fly was stimulated repeatedly with the same odorant stimulus. During one experimental session an equal number of flies were stimulated with the different stimuli (as shown in each data panel) so that day-to-day variation would affect the behavior to all stimuli equally. The order of stimuli was alternated. After each experiment we removed and discarded the fly. Each different experimental paradigm was made up of different sets, depending on the presence and location of the black landing platform. In the experiment shown in Figure 2A, 2B and S3, set 1 placed the landing platform 1.5 cm to the right of the odor source, whereas set 2 place the platform at the odor source directly. In the experiment shown in Figure 2C, S1I and S1J, there was only one set, with the landing platform placed centrally at the location of the odor source. In the experiment shown in Figure 3B and 3D, there was only one set, where the black platform was located 0.5 cm to the right of the odor source. For the experiments shown in Figure 3E, S2A-C, set 1 contained no landing platform, whereas set 2 contained the landing platform at the location of the odor source.

### Differential conditioning

Day 1: Between 15:00 and 16:00, approximately 100 adult flies were removed from standard corn meal agar food and put into a cage (30×30×30 cm, BugDorm-1, BugDorm) that contained a differential conditioning apparatus (Figure 3F). Flies could move around freely at an approximate relative humidity of 30%, a temperature of 25 - 28 °C and normal 12 hour daylight cycle for 24 h.

We trained flies in a differential conditioning paradigm to associate one odorant (positively conditioned stimulus, CS+) with 1 M sucrose solution as the positive reinforcer and to associate another odorant (negatively conditioned stimulus, CS-) with saturated NaCl solution as negative reinforcer (Figure 3F). We used BD and EA as conditioned odorants. We balanced the experiments so that in half of the experiments we used BD as CS+ and EA as CS- and vice versa. CS+ and sucrose solution and CS- and NaCl solution were applied via two horizontally positioned plastic tubes (15 ml, 120 x 17 mm; Sarstedt). Each tube contained 10 ml of either sucrose or NaCl solution and were plugged with a cotton wool to avoid spillage. The frontal 2 cm of each tube remained empty. The odorant was delivered into this empty space via diffusion through a shortened head of a needle (1.2 x 40 mm, Sterican) which ended 1.5 cm inside the empty space of the tube. The needle was connected with a 20 ml glass vial (Schmidlin) that contained the pure odorant and was sealed with a Teflon septum. Thus, to reach the sucrose or NaCl solution, flies had to move through odorized air inside the plastic tube.

Day 2: Between 15:00 and 16:00, the conditioning apparatus was removed and flies were subjected to food and water starvation for the following 24 h in a room with an approximate relative humidity of 60%, a temperature of 25 - 28 °C and normal 12 hour daylight cycle.

Day 3: Flies were tested in the wind tunnel as described above in the section “Experimental protocol for odorants with innate valence” (Day 2).

The conditioning experiments (Figure 3G, S3) also had two sets, depending on the location of the black landing platform. In set 1, the black platform was located 1.5 cm to the right of the odor source, and in set 2, the black platform was at the location of the odor source.

### Stimulus dynamics

To assess the dynamics and precision of the different stimuli, we used a photoionization detector (PID; miniPID model 200B; Aurora Scientific) to record the concentration change of pulses of each of the odorant pairs (BN and BA, BN and BZ, BD and EA) within the wind tunnel. Each pulse had a duration of 500 ms, and an interstimulus interval of 7 s to allow the odorant to clear from the odor delivery device and/or PID and to allow the PID signal to return to baseline before the following pulse was given. We gave a sequence of 100 pulses, alternating between odorant A and odorant B (7 s interval between A and B), thus 50 pulses of each odorant. For each odorant pulse, we calculated the onset time as the time it took to reach 5 % of the maximum PID signal, and the rise time as the time it took for the PID signal to reach from 5 % to 95 % of its maximum. We also calculated the difference in both the onset times and in the rise times between each of the 50 pairs of pulses (A – B).

### Calculating the distance to the target

To calculate the Euclidean distance to the source, we obtained the x, y and z coordinates of the fly for the first 10 s of flight of the recording. For the experiments shown in Fig. 3D, if a fly did not take off, we calculated its closest distance to the target. For all other experiments, if a fly did not take off we took the closest distance between the take-off platform and the target.

For WT 1, we used two cameras which were calibrated within a two pixel scale of each other, thus we did not scale them any further. Both cameras were triggered simultaneously with a TTL pulse, however to ensure that they did not go out of sync, all videos were aligned by first frame of flight. We calculated the Euclidean distance of the fly to the target:

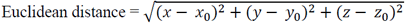

Where x, y and z are the coordinates of the fly’s location in a particular frame, and x0, y0 and z0 are the coordinates of the target.

For WT 2, a single camera was used to film the fly trajectories in the x and y plane. In order to record the movement in the z plane simultaneously, a mirror was placed at 45° to the x-y plane. Thus on the right half of the video recordings, the x-y plane was recorded, and on the left half of the video, the mirrored z-y plane was recorded. However, this led to shrinking of the image in the left half, approximately 1.3 times smaller than the original objects on the right half. Therefore, we calculated the fly’s distance to the target in WT 2 by:
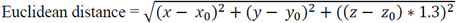

Where x, y and z are the coordinates of the fly’s location in a particular frame, and x0, y0 and z0 are the coordinates of the target

### Quantifying approach with the “half-distance threshold”

In order to measure approach behavior, we used the halfway distance between the frontal border of take-off platform and the target to determine the circular approach area around the target. In WT 1, we used a value of 117 pixels (3.2 cm) for the radius and in WT 2 a value of 71 pixels (2.7 cm).

### Quantifying approach with the “maximized A-B difference threshold”

In order to make the analysis more sensitive for the difference between the approach probabilities for the attractive and the aversive odorants A and B, we defined an approach area that segregated the flies’ approach probability for A (or CS+) and B (or CS-). To determine the radius of this area, we took the Euclidean distance to target for each fly that was exposed to the attractive odorant A (or CS+) alone or the aversive odorant B (or CS-) alone; those flies that encountered mixtures of odorants were not incorporated in this process. The minimum distances were arranged in ascending order, and at each distance, we counted the number of flies from treatment A and treatment B that were included within this threshold distance. Thus for each of these distances, we calculated the difference in approach probabilities by:

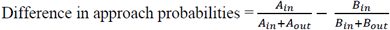

Where *A_in_* represents the number of flies that were presented with odorant A and were included below the threshold, *A_out_* is the number of flies presented with A but excluded above the threshold. *B_in_* and *B_out_* were the same measures for the flies that were presented with odorant B. We then plotted the thresholding index against the vector of minimum distances, and fitted a curve using locally weighted scatterplot smoothing (Figure 3C and S1D, S1E, S2C, S2C, S3B and S3C). We took the distance that corresponded to the maximum peak of the curve as the radius of the approach area, as this point indicates the greatest separation between the two treatment groups. Since we used treatments A and B in defining the approach areas, we did not include these flies in the statistical analyses.

### Approach probability

In both WT 1 and WT 2 we filmed two angles of the flight area. Thus in each wind tunnel, there were two separate areas of approach, one for each of the two cameras for WT 1, and one for each side of the video screen for WT 2 (mirrored and original view). To calculate the approach probability, we gave each fly a binary score. The coordinate of each fly in every frame was recorded and tested as to whether it fell within the approach area boundaries. If a fly entered the approach area at any frame within 10 seconds after take-off, the fly was given a score of 1; if not, was given a score of 0. This was done for each camera (WT 1) or video side (WT 2), and then the results were combined so that only if a fly was in both areas of approach at the same time point, would it be given a score of 1. Finally, we calculated the proportion of flies in each treatment that entered the approach area to get the approach probability.

### Visit probability maps

We extracted the x-y coordinates of the fly during the first ten seconds of flight. We divided the recording image into 20 x 20 pixel bins to create two visit probability maps. Each bin was represented by a cell in the map. We then plotted each coordinate point onto the visit map, giving the cell a score of 1 if one or more points fell into the bin, or a 0 if no points fell into the bin. A matrix of zeros was generated for those flies that did not fly from the entry platform. We calculated the mean for each pixel bin across all of the flies in a treatment group.

### Response latency

We selected the flies that started flying within 10 000 frames after entering the take-off platform (111 s, corresponding to approximately 50 odorant pulses). We defined the individual response latency for each fly as the time point of flight minus the time point of entry onto the take-off platform.

### Statistical Analysis

All statistics were performed using Bayesian data analysis, based on [58]. To compare the approach probabilities, we fitted a generalized linear model using the iteratively reweighted least squares method for fitting. We assumed a flat prior and set a binomial family due to the binary nature of the data: 
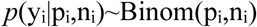

where y_i_ is the number of successes for treatment i, p_i_ is the probability of success for observation i, and n_i_ is the number of trials for treatment i. We used the link function “logit”, which is commonly used for binomial data, to transform the expected values of the outcome variable (probability ranging from 0 to 1) into the range of the linear predictor. We extracted the estimated model parameters for each treatment and then back-transformed the linear predictor to the scale of the outcome variable.

We simulated 100 000 values from the joint posterior distribution of the model parameters. To obtain the fitted value for each treatment, we derived the linear predictor by multiplying the model matrix with the corresponding set of model parameters for each set of simulated values, and then back-transformed the results. We extracted the 2.5 % and the 97.5 % quantiles, creating a 95 % credible interval.

To calculate the certainties that one treatment group had a significantly different approach probability to another group, we compared pairs of treatment groups individually. The proportion of simulations in which one treatment group was higher than that of the compared treatment group represents the posterior probability that the first treatment group has a higher approach probability than the second group. In the figures, we used stars for comparisons between the synchronous mixture AB and the asynchronous mixtures, and we used different letters for comparisons between all stimuli. If the posterior probability was greater than 0.95, we determined the approach probabilities as significantly different (* or different letters). If the posterior probability was greater than 0.99 or 0.999, we indicated their significance as ** and *** respectively (not indicated for comparisons between all stimuli, see text for exact posterior probabilities).

To compare the response latencies across treatment groups, we fitted a linear model using the synchronous mixture AB as the reference level. Similar to an ANOVA, this fits a linear regression to the dataset but using a categorical predictor variable instead of a continuous one. Here, treatment is the categorical variable, which has several indicator variables. AB was always used as the reference level, thus the other indicator variables were either A, B, A33B, and B33A, or A, B, B5A, B10A and B33A, depending on the experimental design. The former is demonstrated in the equation below:

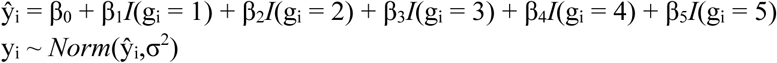

Where y_i_ is the i-th observation and each β value corresponds to the model coefficients for each treatment group g. The residual variance is σ^2^. We simulated from the posterior distribution of the model parameters 100 000 times to obtain the group means and the 2.5 % and 97.5 % quantiles.

To determine whether one treatment group showed a significantly higher response latency compared to another group, we obtained the posterior distribution of the difference between the means of the two groups, by calculating the difference for each draw from the joint posterior distribution of the group means. We then calculated the proportion of draws from the joint posterior distribution for which the mean of the first group was higher than the second group. If the posterior probability was higher than 0.95, it was deemed significantly different (*). If the posterior probability was higher than 0.99 or 0.999, we indicated their significance as ** and *** respectively. For all data analysis, R version 3.5.0 (“Joy in Playing”) were used [59].

## SUPPLEMENTAL INFORMATION

**Table S1.** Percentage of flies flying and their latency to flight.

| Experiment | Stimulus | Total number of flies tested | Percentage of flies flying | Mean latency to flight (s) |
| --- | --- | --- | --- | --- |
| Figure 2B,<br>Figure S1D-H | A (BN) | 42 | 86 | 13 |
|  | B (BZ) | 37 | 68 | 27 |
|  | AB | 39 | 80 | 10 |
|  | A33B | 41 | 90 | 15 |
|  | B33A | 41 | 90 | 16 |
| Figure 2C | A (BN) | 46 | 85 | 10 |
|  | B (BA) | 45 | 76 | 12 |
|  | Air | 41 | 71 | 28 |
| Figure 3A | A (BN) | 93 | 90 | 21 |
|  | B (BA) | 91 | 76 | 16 |
|  | AB | 85 | 72 | 19 |
|  | B33A | 94 | 78 | 20 |
| Figure 3D | A (BN) | 44 | 96 | 19 |
|  | B (BA) | 43 | 84 | 21 |
|  | AB | 41 | 80 | 17 |
|  | B5A | 43 | 88 | 20 |
|  | B10A | 45 | 80 | 21 |
|  | B33A | 45 | 87 | 22 |
| Figure 3E,<br>Figure S2 | A (BN) | 49 | 86 | 21 |
|  | B (BA) | 48 | 70 | 15 |
|  | AB | 44 | 64 | 18 |
|  | A33B | 50 | 72 | 21 |
|  | B33A | 49 | 70 | 17 |
| Figure 3G,<br>Figure S3 | CS+ (EA/BD) | 40 | 88 | 21 |
|  | CS- (BD/EA) | 43 | 86 | 12 |
|  | CS+CS- | 45 | 84 | 17 |
|  | CS+33CS- | 44 | 92 | 7 |
|  | CS-33CS+ | 50 | 98 | 12 |

**Figure S1.**
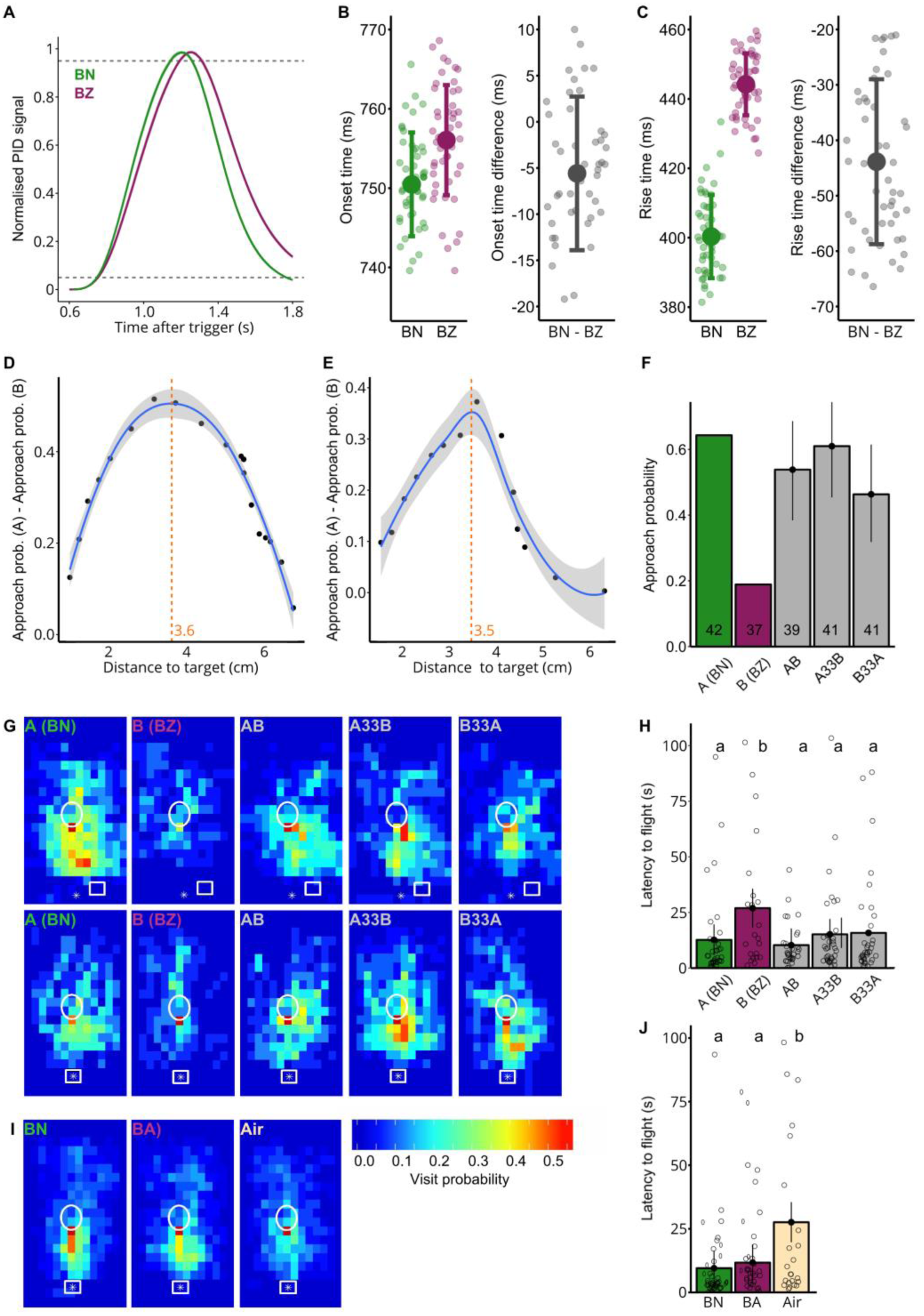
Stimulus timing and behavior data for 2- butanone and benzaldehyde. Related to Figure 2 A, B.(**A**) PID recordings of pulsed stimuli for the odorant pair with innate valence 2- butanone (BN, green) and benzaldehyde (BZ, magenta). Pulses were 500 ms long and with a 7 s interstimulus interval to allow the concentration to reach baseline again before the next pulse started (mean and SD over 50 pulses). Each PID signal was normalized to the maximum concentration reached. Grey dashed lines represent 5 and 95 % of the maximum. (**B**) Left: Onset time (time taken to reach 5 % of maximum concentration after valve trigger) for BN and BZ (mean and SD over 50 pulses). Individual points represent the onsets for each pulse. Right: Onset time difference between pairs of BN and BZ pulses (mean and SD over 50 pulses). (**C**) Left: Rise time (time take to reach 95 % of maximum concentration from the 5% onset time) for BN and BZ (mean and SD over 50 pulses). Individual points represent the rise times for each pulse. Right: Mean rise time difference between pairs of BN and BZ pulses (mean and SD over 50 Pulses). (**D**) Thresholding method that uses the distance which separates flies’ approach probabilities for (BN) A and (BZ) B best for set 1 (see “maximized A-B difference threshold” in Methods). Each point represents the proportion of A-stimulated flies that approached the target by the given minimum distance to the target minus the proportion of B-stimulated flies. The blue trend line was fitted using locally weighted scatterplot smoothing. The distance at the peak of the trend line was defined as threshold (orange dashed line and value) (**E**) Same as (D) but for set 2 of BN and BZ. (**F**) Approach probability for odorant mixtures with different asynchronies (maximized A-B difference threshold). Bars with error bars represent the fitted mean and 95 % credible intervals. Since A and B are used to determine the threshold, they were not included in the statistical analysis and the observed rather than fitted means are shown. (**G**) Visit probability maps of set 1 (Top) and set 2 (Bottom) of BN and BZ for single odorants and the mixtures. The take-off platform (white circle), landing platform (white rectangle) and odor source (white star) are indicated for position reference. n of set 1 = 24, 20, 22, 20 and 22; n of set 2 = 18, 17, 17, 21 and 19 for A, B, AB, A33B and B33A respectively. (**H**) Response latency for odorants and mixtures, measured as the time taken from the fly entering the take-off platform to first flight. The lower case letters represent significantly different responses to the odorant treatments. n = 36, 25, 31, 37 and 37 for A, B, AB, A33B and B33A respectively. (**I**) Same as (G) but for BN (A), BA (B) and blank air control (Air). n = 46, 45 and 41 for A, B, and C respectively. (**J**) Same as (H) but for BN (A), BA (B) and blank air control (Air). n = 39, 34 and 29 for A, B and C respectively.

**Figure S2.**
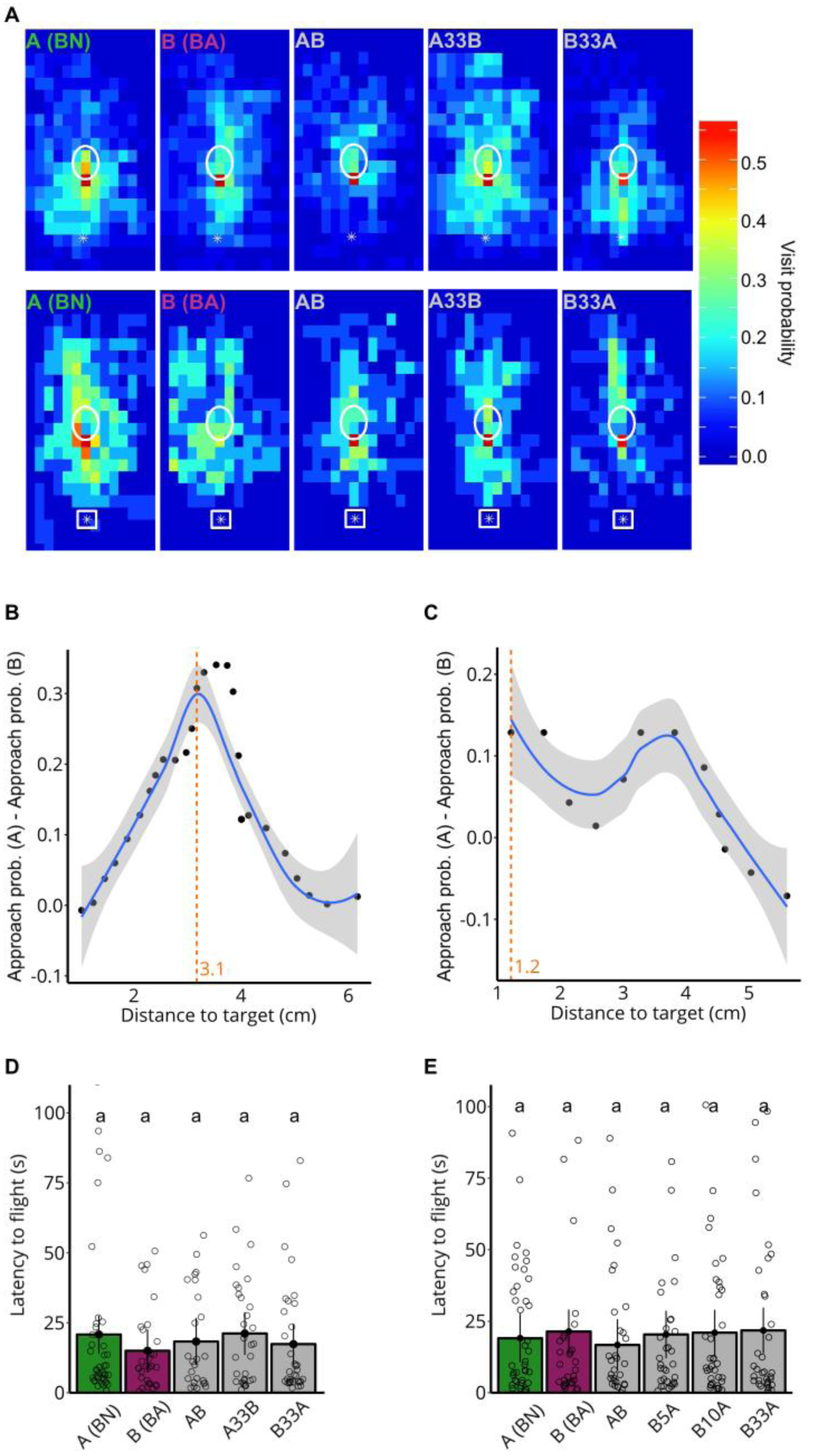
Behavioral data for 2-butanone and butanal. Related to Figure 3D and 3E. (**A**) Visit probability maps of set 1 (Top) and set 2 (Bottom) of BN and BA for single odorants and the mixtures. The take-off platform (white circle), landing platform (white rectangle) and odor source (white star) are indicated for position reference. n for set 1 = 35, 34, 32, 35 and 33; n for set 2 = 14, 14, 12, 15 and 16 for A, B, AB, A33B and B33A respectively. (**B**) Thresholding method that uses the distance which separates flies’ approach probabilities for A and B best for set 1 of BN (A) and BA (B) (maximized A-B difference threshold). (**C**) Same as (B) but for set 2 of BN and BA. (**D**) Response latency for odorants and mixtures, measured as the time taken from the fly entering the take-off platform to first flight for the experiment shown in Figure 3E. The lower case letters represent significantly different responses to the odorant treatment. n = 42, 33, 28, 36 and 34 for A, B, AB, A33B and B33A respectively. (**E**) Same as in (D) but for the experiment shown in Figure 3D. n = 42, 36, 33, 38, 36 and 39 for A, B, AB, B5A, B10A and B33A respectively.

**Figure S3.**
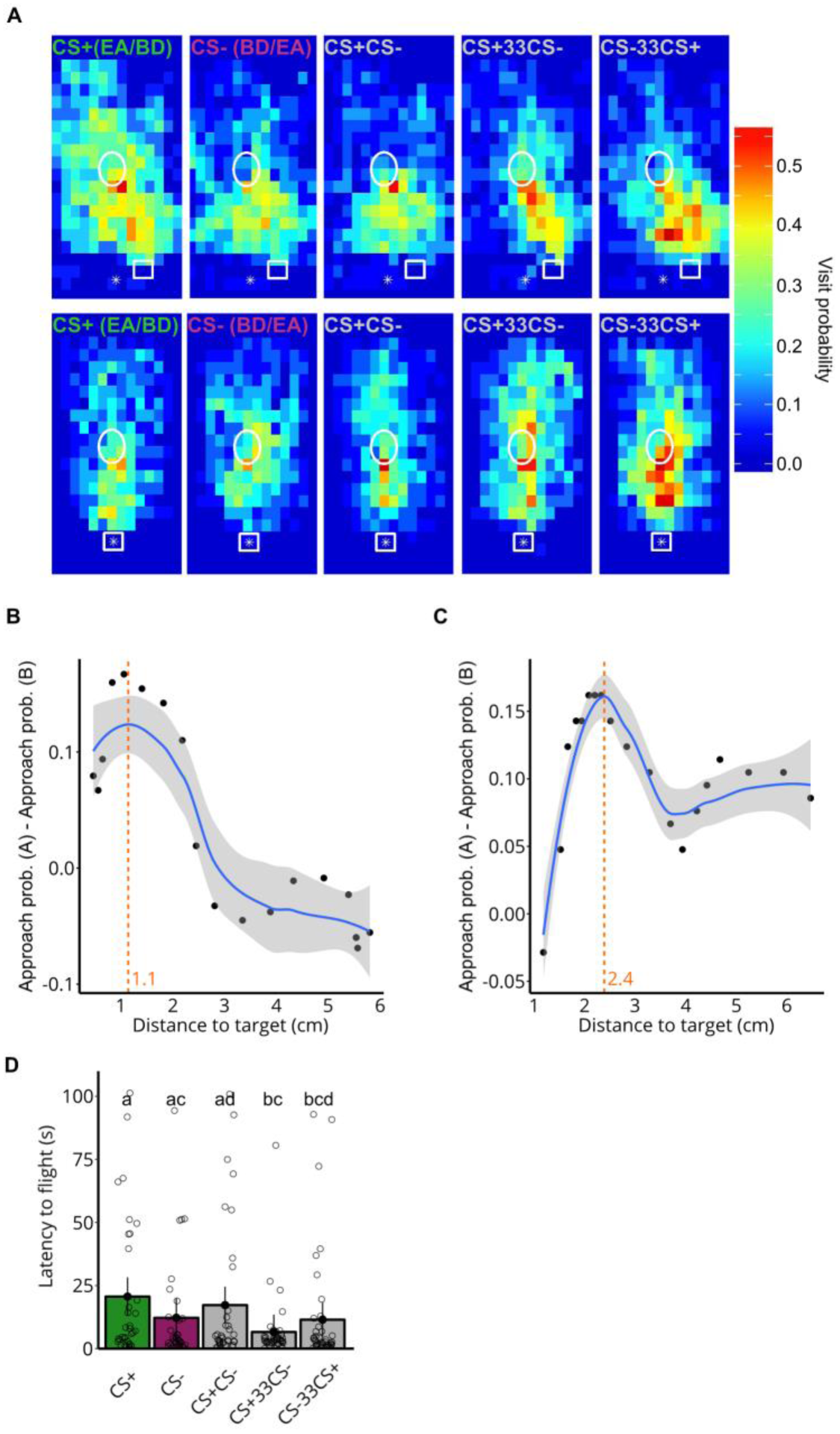
Behavioral data for conditioned odorants. Related to Figure 3G. (**A**) Visit probability maps of set 1 (Top) and set 2 (Bottom) of CS+ and CS- (either EA or BD) for single odorants and the mixtures. The take-off platform (white circle), landing platform (white rectangle) and odor source (white star) are indicated for position reference. n for set 1 = 19, 22, 23, 24 and 26; n for set 2 = 21, 21, 22, 20 and 24 for CS+, CS-, CS+CS-, CS+33CS- and CS-33CS+ respectively. (**B**) Thresholding method that uses the distance which separates flies’ approach probabilities for CS+ and CS-best for set 1 of EA and BD (maximized A-B difference threshold) (**C**) Same as (B) but for set 2 of EA and BD. (**D**) Response latency for odorants and mixtures, measured as the time taken from the fly entering the take-off platform to first flight. The lower case letters represent significantly different responses to the odorant treatments. n = 35, 37, 38, 43 and 46 for CS+, CS-, CS+CS-, CS+33CS- and CS-33CS+ respectively.

## References

1. Kourtzi, Z., and Connor, C.E. (2011). Neural representations for object perception: structure, category, and adaptive coding. Annu. Rev. Neurosci. 34, 45–67. Available at: http://www.annualreviews.org/doi/10.1146/annurev-neuro-060909-153218 [Accessed August 21, 2018].

2. Bizley, J.K., and Cohen, Y.E. (2013). The what, where and how of auditory-object perception. Nat. Rev. Neurosci. 14, 693–707. Available at: http://www.nature.com/doifinder/10.1038/nrn3565 [Accessed August 21, 2018].

3. Galizia, C.G. (2014). Olfactory coding in the insect brain: Data and conjectures. Eur. J. Neurosci. 39, 1784–1795. Available at: http://doi.wiley.com/10.1111/ejn.12558 [Accessed March 25, 2017].

4. Uchida, N., Poo, C., and Haddad, R. (2014). Coding and Transformations in the Olfactory System. Annu. Rev. Neurosci. 37, 363–385. Available at: http://www.ncbi.nlm.nih.gov/pubmed/24905594 [Accessed January 17, 2018].

5. Hopfield, J.J. (1991). Olfactory computation and object perception. Proc. Natl. Acad. Sci. 88, 6462–6466. Available at: http://www.pnas.org/content/88/15/6462.short [Accessed February 17, 2016].

6. Celani, A., Villermaux, E., and Vergassola, M. (2014). Odor landscapes in turbulent environments. Phys. Rev. X 4, 1–17.

7. Hopfield, J.F., and Gelperin, A. (1989). Differential conditioning to a compound stimulus and its components in the terrestrial mollusc Limax maximus. Behav. Neurosci. 103, 329–333. Available at: http://doi.apa.org/getdoi.cfm?doi=10.1037/0735-7044.103.2.329 [Accessed February 17, 2016].

8. Baker, T.C., Fadamiro, H.Y., and Cosse, A.A. (1998). Moth uses fine tuning for odour resolution. Nature 393, 530–530. Available at: http://dx.doi.org/10.1038/31131 [Accessed February 17, 2016].

9. Andersson, M.N., Binyameen, M., Sadek, M.M., and Schlyter, F. (2011). Attraction Modulated by Spacing of Pheromone Components and Anti-attractants in a Bark Beetle and a Moth. J. Chem. Ecol. 37, 899–911. Available at: http://www.ncbi.nlm.nih.gov/pubmed/21750948 [Accessed September 26, 2011].

10. Szyszka, P., Stierle, J.S.J.S., Biergans, S., and Galizia, C.G.G. (2012). The speed of smell: odor-object segregation within milliseconds. PLoS One 7, e36096. Available at: http://www.pubmedcentral.nih.gov/articlerender.fcgi?artid=3338635&tool=pmcentrez&rendertype=abstract [Accessed November 5, 2012].

11. Saha, D., Leong, K., Li, C., Peterson, S., Siegel, G., and Raman, B. (2013). A spatiotemporal coding mechanism for background-invariant odor recognition. Nat. Neurosci. 16, 1830–9. Available at: http://www.ncbi.nlm.nih.gov/pubmed/24185426.

12. Weissburg, M., Atkins, L., Berkenkamp, K., and Mankin, D. (2012). Dine or dash? Turbulence inhibits blue crab navigation in attractive-aversive odor plumes by altering signal structure encoded by the olfactory pathway. J. Exp. Biol. 215, 4175–82. Available at: http://www.ncbi.nlm.nih.gov/pubmed/23136153.

13. Badel, L., Ohta, K., Tsuchimoto, Y., and Kazama, H. (2016). Decoding of Context-Dependent Olfactory Behavior in Drosophila. Neuron 91, 155–167. Available at: http://dx.doi.org/10.1016/j.neuron.2016.05.022.

14. Martelli, C., Carlson, J.R., and Emonet, T. (2013). Intensity invariant dynamics and odor-specific latencies in olfactory receptor neuron response. J. Neurosci. 33, 6285–6297. Available at: http://www.jneurosci.org/cgi/doi/10.1523/JNEUROSCI.0426-12.2013 [Accessed June 4, 2013].

15. Raiser, G., Galizia, C.G.G., and Szyszka, P. (2016). A High-Bandwidth Dual-Channel Olfactory Stimulator for Studying Temporal Sensitivity of Olfactory Processing. Chem. Senses 42, bjw114. Available at: http://chemse.oxfordjournals.org/lookup/doi/10.1093/chemse/bjw114 [Accessed March 25, 2017].

16. Budick, S.A., and Dickinson, M.H. (2006). Free-flight responses of Drosophila melanogaster to attractive odors. J. Exp. Biol. 209, 3001–3017. Available at: http://www.ncbi.nlm.nih.gov/pubmed/16857884 [Accessed August 21, 2018].

17. Houot, B., Gigot, V., Robichon, A., and Ferveur, J.-F. (2017). Free flight odor tracking in Drosophila: Effect of wing chemosensors, sex and pheromonal gene regulation. Sci. Rep. 7, 40221. Available at: http://www.nature.com/articles/srep40221 [Accessed August 21, 2018].

18. Saxena, N., Natesan, D., and Sane, S.P. (2018). Odor source localization in complex visual environments by fruit flies. J. Exp. Biol. 221, jeb172023. Available at: http://www.ncbi.nlm.nih.gov/pubmed/29146771 [Accessed January 13, 2018].

19. Breugel, F. van, Huda, A., and Dickinson, M.H. (2017). Drosophila have distinct activity-gated pathways that mediate attraction and aversion to CO2. bioRxiv, 227991. Available at: https://www.biorxiv.org/content/early/2017/12/03/227991 [Accessed August 23, 2018].

20. Gorea, A. (1987). Masking efficiency as a function of stimulus onset asynchrony for spatial-frequency detection and identification. Spat. Vis. 2, 51–60. Available at: http://www.ncbi.nlm.nih.gov/pubmed/3154937 [Accessed October 13, 2018].

21. Tully, T. (1984). Drosophila learning: Behavior and biochemistry. Behav. Genet. 14, 527–557. Available at: http://uml.idm.oclc.org/login?url=http://search.proquest.com/docview/617053337?accountid=14569%5Cn http://primo-pmtna01.hosted.exlibrisgroup.com/openurl/01UMB_INST/umb_services_page??url_ver=Z39.88-2004&rft_val_fmt=info:ofi/fmt:kev:mtx:journal&genre=artic [Accessed March 25, 2017].

22. Nikonov, A.A., and Leal, W.S. (2002). Peripheral coding of sex pheromone and a behavioral antagonist in the Japanese beetle, Popillia japonica. J. Chem. Ecol. 28, 1075–89. Available at: http://www.ncbi.nlm.nih.gov/pubmed/12049228.

23. Hukin, R.W., and Darwin, C.J. (1995). Comparison of the effect of onset asynchrony on auditory grouping in pitch matching and vowel identification. Percept. Psychophys. 57, 191–6. Available at: http://www.ncbi.nlm.nih.gov/pubmed/7885817 [Accessed October 12, 2018].

24. Lipp, R., Kitterick, P., Summerfield, Q., Bailey, P.J., and Paul-Jordanov, I. (2010). Concurrent sound segregation based on inharmonicity and onset asynchrony. Neuropsychologia 48, 1417–25. Available at: http://linkinghub.elsevier.com/retrieve/pii/S0028393210000102 [Accessed October 12, 2018].

25. Usher, M., and Donnelly, N. (1998). Visual synchrony affects binding and segmentation in perception. Nature 394, 179–182. Available at: http://www.ncbi.nlm.nih.gov/pubmed/9671300 [Accessed August 23, 2018].

26. Hancock, P.J.B., Walton, L., Mitchell, G., Plenderleith, Y., and Phillips, W.A. (2008). Segregation by onset asynchrony. J. Vis. 8, 21. Available at: http://jov.arvojournals.org/article.aspx?doi=10.1167/8.7.21 [Accessed October 12, 2018].

27. Jefferis, G.S.X.E., Potter, C.J., Chan, A.M., Marin, E.C., Rohlfing, T., Maurer, C.R., and Luo, L. (2007). Comprehensive Maps of Drosophila Higher Olfactory Centers: Spatially Segregated Fruit and Pheromone Representation. Cell 128, 1187–1203. Available at: http://www.ncbi.nlm.nih.gov/pubmed/17382886 [Accessed August 22, 2018].

28. Roussel, E., Carcaud, J., Combe, M., Giurfa, M., and Sandoz, J.-C. (2014). Olfactory Coding in the Honeybee Lateral Horn. Curr. Biol. 24, 561–567. Available at: http://www.ncbi.nlm.nih.gov/pubmed/24560579 [Accessed August 22, 2018].

29. Strutz, A., Soelter, J., Baschwitz, A., Farhan, A., Grabe, V., Rybak, J., Knaden, M., Schmuker, M., Hansson, B.S., and Sachse, S. (2014). Decoding odor quality and intensity in the Drosophila brain. Elife 3, e04147. Available at: http://www.ncbi.nlm.nih.gov/pubmed/25512254 [Accessed March 27, 2018].

30. Jeanne, J.M., Fişek, M., and Wilson, R.I. (2018). The Organization of Projections from Olfactory Glomeruli onto Higher-Order Neurons. Neuron 98, 1198–1213.e6. Available at: http://www.ncbi.nlm.nih.gov/pubmed/29909998 [Accessed August 22, 2018].

31. Strube-Bloss, M.F., Nawrot, M.P., and Menzel, R. (2011). Mushroom Body Output Neurons Encode Odor – Reward Associations. J. Neurosci. 31, 3129–3140. Available at: http://www.jneurosci.org/cgi/doi/10.1523/JNEUROSCI.2583-10.2011.

32. Aso, Y., Sitaraman, D., Ichinose, T., Kaun, K.R., Vogt, K., Belliart-Guérin, G., Plaçais, P.Y., Robie, A.A., Yamagata, N., Schnaitmann, C., et al. (2014). Mushroom body output neurons encode valence and guide memory-based action selection in Drosophila. Elife 3, e04580. Available at: http://elifesciences.org/lookup/doi/10.7554/eLife.04580 [Accessed March 25, 2017].

33. Hige, T., Aso, Y., Modi, M.N., Rubin, G.M., and Turner, G.C. (2015). Heterosynaptic Plasticity Underlies Aversive Olfactory Learning in Drosophila. Neuron 88, 985–998. Available at: http://linkinghub.elsevier.com/retrieve/pii/S0896627315009824 [Accessed March 25, 2017].

34. Smith, B.H. (1998). Analysis of interaction in binary odorant mixtures. Physiol. Behav. 65, 397–407. Available at: http://www.ncbi.nlm.nih.gov/pubmed/9877404 [Accessed August 22, 2018].

35. Chandra, S., and Smith, B.H. (1998). An analysis of synthetic processing of odor mixtures in the honeybee (Apis mellifera). J. Exp. Biol. 201, 3113–21. Available at: http://www.ncbi.nlm.nih.gov/pubmed/9787131 [Accessed August 22, 2018].

36. Deisig, N., Lachnit, H., Giurfa, M., and Hellstern, F. (2001). Configural olfactory learning in honeybees: Negative and positive patterning discrimination. Learn. Mem. 8, 70–78. Available at: http://www.ncbi.nlm.nih.gov/pubmed/11274252 [Accessed August 22, 2018].

37. Broome, B.M., Jayaraman, V., and Laurent, G. (2006). Encoding and Decoding of Overlapping Odor Sequences. Neuron 51, 467–482. Available at: http://www.ncbi.nlm.nih.gov/pubmed/16908412 [Accessed August 22, 2018].

38. Deisig, N., Giurfa, M., Lachnit, H., and Sandoz, J.C. (2006). Neural representation of olfactory mixtures in the honeybee antennal lobe. Eur. J. Neurosci. 24, 1161–1174. Available at: http://www.ncbi.nlm.nih.gov/pubmed/16930442.

39. Silbering, A.F., and Galizia, C.G. (2007). Processing of odor mixtures in the Drosophila antennal lobe reveals both global inhibition and glomerulus-specific interactions. J. Neurosci. 27, 11966–77. Available at: http://dx.doi.org/10.1523/JNEUROSCI.3099-07.2007 [Accessed March 25, 2017].

40. Münch, D., Schmeichel, B., Silbering, A.F., and Galizia, C.G. (2013). Weaker ligands can dominate an odor blend due to syntopic interactions. Chem. Senses 38, 293–304. Available at: http://www.ncbi.nlm.nih.gov/pubmed/23315042 [Accessed March 25, 2017].

41. Münch, D., and Galizia, C.G. (2017). Take time: odor coding capacity across sensory neurons increases over time in Drosophila. J. Comp. Physiol. A 203, 959–972. Available at: http://www.ncbi.nlm.nih.gov/pubmed/28852844 [Accessed September 10, 2018].

42. Nowotny, T., Stierle, J.S.J.S., Galizia, C.G.G., and Szyszka, P. (2013). Data-driven honeybee antennal lobe model suggests how stimulus-onset asynchrony can aid odour segregation. Brain Res. 1536, 119–34. Available at: http://dx.doi.org/10.1016/j.brainres.2013.05.038 [Accessed March 25, 2017].

43. Stierle, J.S.J.S., Giovanni Galizia, C., Szyszka, P., Galizia, C.G., and Szyszka, P. (2013). Millisecond Stimulus Onset-Asynchrony Enhances Information about Components in an Odor Mixture. J. Neurosci. 33, 6060–6069. Available at: http://www.jneurosci.org/cgi/doi/10.1523/JNEUROSCI.5838-12.2013%5Cnpapers3://publication/doi/10.1523/JNEUROSCI.5838-12.2013 [Accessed March 25, 2017].

44. Laloi, D., Roger, B., Blight, M.M., Wadhams, L.J., and Pham-Delegue, M.H. (1999). Individual learning ability and complex odor recognition in the honey bee, Apis mellifera L. J. Insect Behav. 12, 585–597.

45. Reinhard, J., Sinclair, M., Srinivasan, M. V., and Claudianos, C. (2010). Honeybees Learn Odour Mixtures via a Selection of Key Odorants. PLoS One 5, e9110. Available at: http://www.ncbi.nlm.nih.gov/pubmed/20161714 [Accessed August 22, 2018].

46. Smith, B.H., and Cobey, S. (1994). The olfactory memory of the honeybee Apis mellifera. II. Blocking between odorants in binary mixtures. J. Exp. Biol. 195, 91–108. Available at: http://www.ncbi.nlm.nih.gov/pubmed/7964421 [Accessed May 14, 2017].

47. Thoma, M., Hansson, B.S., and Knaden, M. (2014). Compound valence is conserved in binary odor mixtures in Drosophila melanogaster. J. Exp. Biol. 217, 3645–3655. Available at: http://www.ncbi.nlm.nih.gov/pubmed/25189369 [Accessed August 23, 2018].

48. Young, J.M., Wessnitzer, J., Armstrong, J.D., and Webb, B. (2011). Elemental and non-elemental olfactory learning in Drosophila. Neurobiol. Learn. Mem. 96, 339–352. Available at: http://www.ncbi.nlm.nih.gov/pubmed/21742045 [Accessed August 22, 2018].

49. Campbell, R.A.A., Honegger, K.S., Qin, H., Li, W., Demir, E., and Turner, G.C. (2013). Imaging a population code for odor identity in the Drosophila mushroom body. J. Neurosci. 33, 10568–81. Available at: http://www.jneurosci.org/content/33/25/10568.abstract.

50. Mayer, M.L., Westbrook, G.L., and Guthrie, P.B. (1984). Voltage-dependent block by Mg2+of NMDA responses in spinal cord neurones. Nature 309, 261–263. Available at: http://www.ncbi.nlm.nih.gov/pubmed/6325946 [Accessed August 24, 2018].

51. Sinakevitch, I., Grau, Y., Strausfeld, N.J., and Birman, S. (2010). Dynamics of glutamatergic signaling in the mushroom body of young adult Drosophila. Neural Dev. 5, 10. Available at: http://neuraldevelopment.biomedcentral.com/articles/10.1186/1749-8104-5-10 [Accessed August 24, 2018].

52. Owald, D., Felsenberg, J., Talbot, C.B., Das, G., Perisse, E., Huetteroth, W., and Waddell, S. (2015). Activity of defined mushroom body output neurons underlies learned olfactory behavior in Drosophila. Neuron 86, 417–427. Available at: http://www.ncbi.nlm.nih.gov/pubmed/25864636 [Accessed February 14, 2017].

53. Sato, K., Pellegrino, M., Nakagawa, T., Nakagawa, T., Vosshall, L.B., and Touhara, K. (2008). Insect olfactory receptors are heteromeric ligand-gated ion channels. Nature 452, 1002–1006.

54. Schuckel, J., Meisner, S., Torkkeli, P.H., and French, A.S. (2008). Dynamic properties of Drosophila olfactory electroantennograms. J. Comp. Physiol. A Neuroethol. Sensory, Neural, Behav. Physiol. 194, 483–489. Available at: http://link.springer.com/10.1007/s10886-011-9995-3.

55. Szyszka, P., Gerkin, R.C., Galizia, C.G.G., and Smith, B.H.B.H. (2014). High-speed odor transduction and pulse tracking by insect olfactory receptor neurons. Proc. Natl. Acad. Sci. U. S. A. 111, 16925–30. Available at: http://www.pubmedcentral.nih.gov/articlerender.fcgi?artid=4250155&tool=pmcentrez&rendertype=abstract%5Cnhttp://www.ncbi.nlm.nih.gov/pubmed/25385618%5Cnhttp://www.pubmedcentral.nih.gov/articlerender.fcgi?artid=PMC4250155 [Accessed March 25, 2017].

56. Egea-Weiss, A., Renner, A., Kleineidam, C.J., and Szyszka, P. (2018). High Precision of Spike Timing across Olfactory Receptor Neurons Allows Rapid Odor Coding in Drosophila. iScience 4, 76–83. Available at: https://linkinghub.elsevier.com/retrieve/pii/S2589004218300646 [Accessed August 22, 2018].

57. Krofczik, S., Menzel, R., and Nawrot, M.P. (2009). Rapid odor processing in the honeybee antennal lobe network. Front. Comput. Neurosci. 2, 9. Available at: http://www.ncbi.nlm.nih.gov/pubmed/19221584 [Accessed March 25, 2017].

58. Korner-Nievergelt, F., Roth, T., von Felten, S., Guélat, J., Almasi, B., and Korner-Nievergelt, P. (2015). Bayesian data analysis in ecology using linear models with R, BUGS, and Stan (Academic Press).

59. R Core Team (2012). R: A Language and Environment for Statistical Computing. R Foundation for Statistical Computing. R Found. Stat. Comput. Vienna, Austria. Available at: http://www.r-project.org [Accessed August 27, 2018].

